# TandemAligner: a new parameter-free framework for fast sequence alignment

**DOI:** 10.1101/2022.09.15.507041

**Authors:** Andrey V. Bzikadze, Pavel A. Pevzner

**Affiliations:** Graduate Program in Bioinformatics and Systems Biology, University of California, San Diego, CA, USA; Department of Computer Science and Engineering, University of California, San Diego, CA, USA

## Abstract

The recent advances in “complete genomics” revealed the previously inaccessible genomic regions (such as centromeres) and enabled analysis of their associations with diseases. However, analysis of variations in centromeres, immunoglobulin loci, and other extra-long tandem repeats (ETRs) faces an algorithmic bottleneck since there are currently no tools for accurate sequence comparison of ETRs. Counterintuitively, the classical alignment approaches, such as the Smith-Waterman algorithm, that work well for most sequences, fail to construct biologically adequate alignments of ETRs. This limitation was overlooked in previous studies since the ETR sequences across multiple genomes only became available in the last year. We present TandemAligner — the first parameter-free sequence alignment algorithm that introduces a sequence-dependent alignment scoring that automatically changes for any pair of compared sequences. We apply TandemAligner to various human centromeres and primate immunoglobulin loci, arrive at the first accurate estimate of the mutation rates in human centromeres, and quantify the extremely high rate of large insertions/duplications in centromeres. This extremely high rate (that the standard alignment algorithms fail to uncover) suggests that centromeres represent the most rapidly evolving regions of the human genome with respect to their structural organization.

## Introduction

The Telomere-to-Telomere Consortium (T2T) recently assembled the first complete sequence of a human genome (Nurk et al. 2022) and the efforts are currently underway to generate hundreds of complete haplotype-resolved human genomes (Wang et al. 2022; Liao et al. 2022). This progress opens a possibility to address the long-standing questions about the variation and structure of many biomedically important and complex regions of the genome such as segmental duplications (Vollger, Guitart, et al. 2022) and long tandem repeats (Hoyt et al. 2022). Since tandem repeats rapidly accumulate changes in the copy numbers, they greatly differ across the human population and their expansion is known to be the causal factor of various diseases (Bakhtiari et al. 2021; Park et al. 2022). For example, alpha satellite arrays that host centromeres are formed by *extra-long tandem repeats* (*ETRs*) representing some of the most rapidly evolving regions of the human genome (Dvorkina et al. 2021; Altemose et al. 2022; Kunyavskaya et al. 2022). Comparison of ETRs across the human population is a prerequisite for understanding their evolution and association with cancer and infertility (Schueler et al. 2001; Alkan et al. 2007; Enukashvily et al. 2007; Shepelev et al. 2009; Ting et al. 2011; Nagaoka, Hassold, and Hunt 2012; Melters et al. 2013; Giunta and Funabiki 2017; Black and Giunta 2018; Smurova and De Wulf 2018; Miga 2019; Miga and Alexandrov 2021; Altemose et al. 2022). However, alignment of ETRs across various human genomes remains an open algorithmic challenge. As a result, constructing the pancentromere represents a bottleneck for the ongoing effort to generate the human pangenome graph, e.g., centromeres and other ETRs have been excluded from the pangenome graph recently constructed by the Human Pangenome Reference (HPR) consortium (Liao et al. 2022). Comparing the immunoglobulin loci across vertebrate species represents yet another bottleneck since the standard alignment approach fails to adequately align these biomedically important ETRs (Sirupurapu, Safonova, and Pevzner 2022).

A common approach to variant calling is based on aligning reads that originated from the query genome against the target sequence (Xu 2018) to genotype single nucleotide variants (Van der Auwera and O’Connor 2020) and short tandem repeats (Bakhtiari et al. 2018; Park et al. 2022; Mousavi et al. 2019). Emergence of long-read sequencing technologies enabled further genotyping of structural variants that are located outside ETRs (Chaisson et al. 2019) but genotyping ETRs remains an unsolved problem even with long-reads (Liao et al. 2022). Thus, *de novo* genome assembly using long reads is currently the only approach that generates accurate ETR sequences (Bzikadze and Pevzner 2020; Nurk et al. 2020; Cheng et al. 2021; Bankevich et al. 2022; Rautiainen et al. 2022).

Emergence of “complete genomics” (Nurk et al. 2022) and “complete metagenomics” (Bickhart et al. 2022) is shifting the focus of read mapping from variant calling to validation of newly generated assemblies (Mikheenko et al. 2020). Given assembled ETRs, the difficult problem of finding variations in ETRs by read mapping is substituted by a seemingly simpler problem of aligning ETR sequences using the standard dynamic programming approaches such as the Smith-Waterman algorithm (T. F. Smith and Waterman 1981). However, we show that it is an inadequate computational model for analyzing ETRs since the highest-scoring pairwise alignment fails to reveal the evolutionary events that made two ETRs different (for any choice of scoring parameters).

The HPR Consortium recently arrived at a similar conclusion and stated that “*more work is needed to determine how best to align and represent these large repeat arrays within pangenomes, particularly as T2T assembly becomes commonplace*… *new methods may need to be developed to fully understand and characterize this component of the human pangenome*.” (Liao et al. 2022).

We describe the TandemAligner algorithm that addresses the ETR comparison problem and illustrate its performance using human centromeres and primate immunoglobulin loci. We classify a substring of a string *S* as rare if its count in *S* does not exceed a small threshold, and frequent, otherwise. ETRs are rich with frequent substrings, e.g., most 20-mers in the human centromere X occur over a thousand times in this centromere (Miga et al. 2020; Nurk et al. 2022; Altemose et al. 2022). Thus, deciding whether a match between two occurrences of a frequent substring in two ETRs is challenging as the total number of such matches is often measured in millions.

The standard alignment algorithms apply the same scoring parameters (match/mismatch/indel penalty, affine gap penalties, etc.) to *all* pairs of sequences. Although they construct a biologically adequate alignment for most sequences (wrt to their evolution), below we show that some sequences (e.g., ETRs) are less amenable to this approach for any choice of alignment parameters. The inherent limitation of the standard alignment approach is that less significant matches of frequent *k*-mers (that may have millions of matches in an ETR) have the same contribution to the score as the more significant matches of rare *k*-mers (that may have a single match in an ETR). This limitation was overlooked in the previous studies since the sequences of the assembled centromeres (and other ETRs) across multiple genomes only became available in the last year.

TandemAligner introduces a novel *parameter-free* and *sequence-dependent* alignment scoring that automatically changes for any pair of compared sequences and that does not rely on the multiple scoring parameters used in the standard alignment. The sequence-dependent alignment prioritizes matches of rare substrings since they are more likely to be relevant to the evolutionary relationship between two sequences. Analysis of the constructed *rare-alignments* reveals new biological phenomena such as the extremely high rate of large insertion/deletion in centromeres. Since the T2T Consortium did not have bioinformatics tools for deriving the detailed history of indels in centromeres, its analysis of the first two assembled human centromeres resulted in a conclusion that they are highly concordant apart from the three regions with recent large indels (Figure 5D in (Altemose et al. 2022)). We show that these centromeres differ from each other by over 300 large deletions and duplications of length at least 2 kb, including six extra-long indels varying in length from 23 to 76 kb. This extremely high rate of large duplications and deletions (that the standard alignment algorithms fail to uncover) suggests that centromeres represent the most rapidly evolving regions of the human genome with respect to their structural organization.

Analysis of rare-alignments also leads to the first accurate estimate of the single-nucleotide mutation rates in human centromeres. Previous studies came to the conclusion that the rate of single-nucleotide mutations is greatly elevated in centromeres (Rudd et al. 2006; Pertile et al. 2009). However, in the absence of bioinformatics tools for accurate ETR alignments, it is nearly impossible to identify orthologous repeat copies, a prerequisite for estimating single-nucleotide mutation rates. Rare-alignments revealed that, contrary to existing assumption, the rate of single-nucleotide mutations (and small indels) in centromeres does not exceed the average mutation rate of the human genome.

We demonstrate that TandemAligner not just improves on the standard alignment approach with respect to comparing ETRs (human centromeres and primate immunoglobulin loci) - instead it often generates completely different alignments. However, although TandemAligner was developed for comparing highly-repetitive sequences, we show that it results in fast and accurate comparison of any sequences with percent identity exceeding 70% and even slightly outperforms the standard alignment approach (with respect to accuracy) in the case of highly similar sequences with percent identity exceeding 90%. We further discuss theoretical properties of the alignment problem and suggest that the scoring scheme should be selected with respect to the underlying evolutionary model that is presumed to have produced the aligned sequences. Although the standard scoring scheme for sequence alignment often performs well in practice, it makes some implicit assumptions that do not hold for ETRs. While the rare-alignment scheme that is used by TandemAligner is not necessarily induced by any natural evolutionary model, rare-alignments of centromeres and immunoglobulin loci appear to be closer to the “real alignment” than the traditional alignments.

## Results

### Outline of the TandemAligner algorithm

Figure Algorithm illustrates some (but not all) steps of TandemAligner. The codebase of TandemAligner is available at https://github.com/seryrzu/tandem_aligner.

A substring of a string *S* is called *unique* if it appears exactly once in *S*. TandemAligner uses the *Longest Common Prefix* array (Manber and Myers 1989) to find the *shortest unique* substring starting at each position in the string *S*. A shortest unique substring *P* of a string *S* is called an *anchor* if all proper substrings of *P* are non-unique in *S*. Given strings *S* and *T*, TandemAligner efficiently computes the set of all anchors shared by *S* and *T* (this set is usually small) and uses *sparse dynamic programming* to rapidly construct an optimal path through these anchors in the standard grid-like sequence alignment graph. It further analyzes the resulting alignment-path *Alignment* formed by diagonal edges representing anchors and horizontal/vertical indel edges. We refer to a segment of *Alignment* consisting of a deletion-run immediately followed by an insertion-run (or vice versa) as an *indel-pair*. Each indel-pair corresponds to a pair of substrings *s* and *t* in *S* and *T*, respectively. TandemAligner recursively constructs an alignment of strings *s* and *t*, and substitutes the indel-pair in *Alignment* by the resulting (small) alignment-path (this recursive step is not illustrated in Figure Algorithm). The process terminates when there are either no indel-pairs left or the set of anchors constructed on substrings corresponding to each remaining indel-pair is empty.

**Figure Algorithm.**
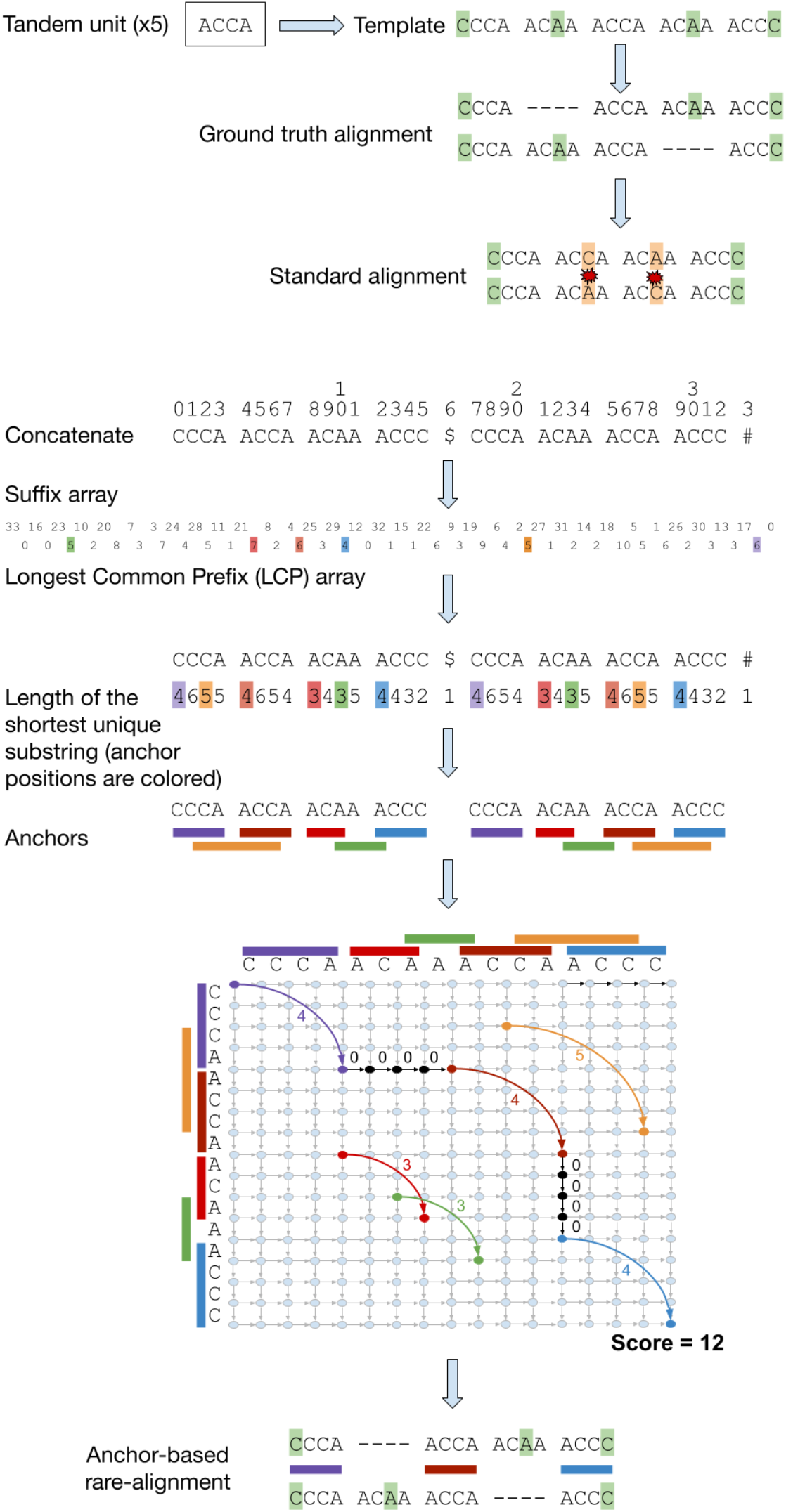
Aligning ETRs *S*=CCCAACCAACAAACCC and *T*=CCCAACAAACCAACCC using TandemAligner. These four-unit ETRs evolved from a five-unit ancestral ETR CCCAACAAACCAACAAACCC by deletion of its second unit in *S* and its fourth unit in *T*. The standard alignment generates high-scoring but incorrect alignment between *S* and *T*. TandemAligner constructs the suffix array and the Longest Common Prefix (LCP) array of the concatenate *S*T* (with delimiters “#” and “$”) and uses these arrays to rapidly identify shortest unique substrings and anchors. Afterward, it constructs a (sparse) anchor graph and finds an optimal alignment-path in this graph using sparse dynamic programming. Finally, it recursively applies the same procedure to substrings of *S* and *T* that form a pair of consecutive insertion-deletion or deletion-insertion edges in the constructed alignment-path (this step is not shown).

### Sequence architecture of human centromeres

Alpha satellite repeats are the building blocks of centromeric repeats and form roughly 3% of the human genome (Altemose et al. 2022). Consecutive alpha satellite repeats are arranged into *higher-order repeat* (*HOR*) units that are repeated hundreds or thousands of times in each centromere. Individual alpha satellites within a higher-order unit show low (50–90%) sequence identity to each other while HORs within a single centromere show high (95–100%) sequence identity (Figure CentromereArchitecture). Organization and nucleotide sequence of HORs is specific for a particular chromosome.

**Figure CentromereArchitecture.**
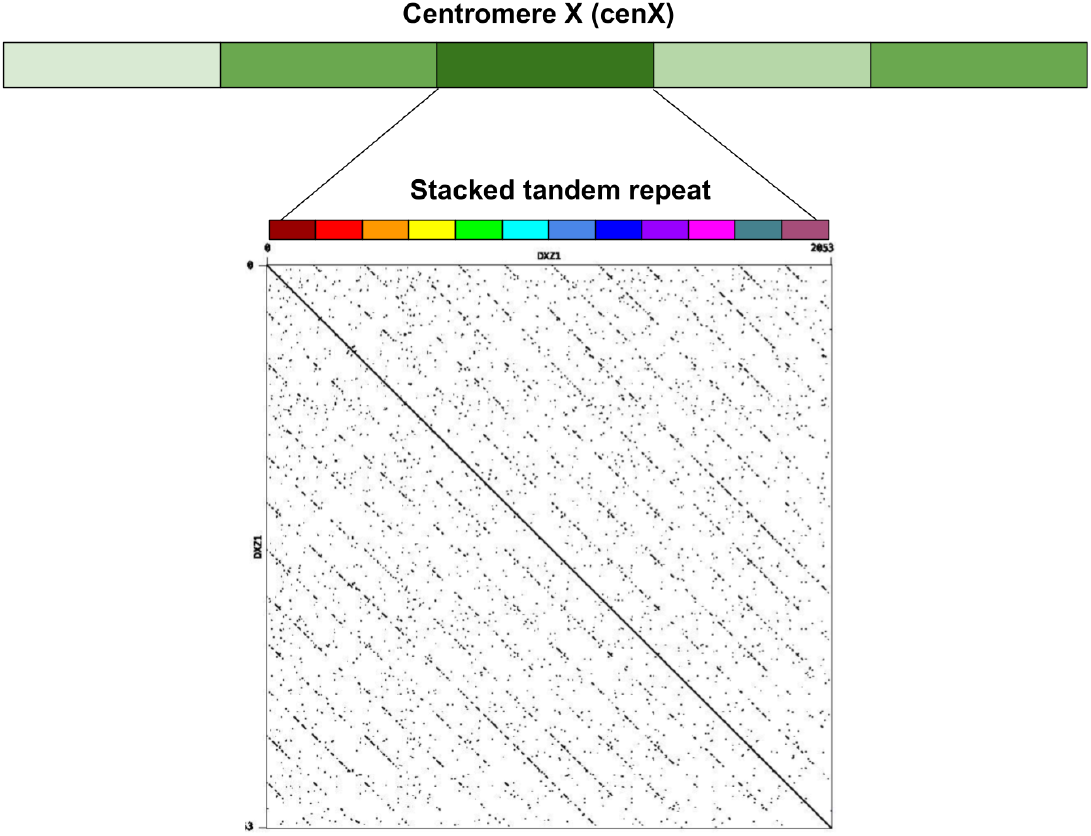
The architecture of centromere on Chromosome X. The centromere of human chromosome X (cenX) consists of ∼18100 monomers of length ≅171 bp each based on the cenX assembly in (Bzikadze and Pevzner 2020) (the T2T assembly (Nurk et al. 2022) represents a minor change to this assembly). These monomers are organized into ∼1500 units. Five units in the Figure are colored by five shades of green illustrating unit variations. Each unit is a stacked tandem repeat formed by various monomers. The vast majority of units in cenX correspond to the canonical HOR which is formed by twelve monomers (shown by twelve different colors). The figure on top represents the dot plot of the nucleotide sequence of the canonical HOR that reveals twelve monomers. While the canonical units are 95–100% similar, monomers in cenX are only 65–88% similar. In addition to the canonical 12-monomer units, cenX has a small number of partial HORs with varying numbers of monomers.

### Datasets

We extracted the assembled satellite centromere on chromosome X (cenX) from the public release v2.0 by the Telomere-to-Telomere (T2T) consortium of the complete assembly for the effectively haploid CHM13 cell line (GCA_009914755.4), (Miga et al. 2020; Logsdon et al. 2021; Nurk et al. 2022). Specifically, we use the following coordinates: chrX:57,817,899-60,927,196. We also extracted cenX from the recently assembled HG002 human sample (CP086568.2, coordinates: chrX:57,860,000-61,000,000). We refer to these two centromeres as *cenX*_1_ and *cenX*_2_. *cenX*_1_ of total length 3,109,297 bp is formed by 1533 HORs and *cenX*_*2*_ of total length 3,140,000 bp is formed by 1539 HORs.

Similar lengths of *cenX*_1_ and *cenX*_2_ may create a wrong impression that the lengths of centromeres is conserved across the human population. It is known that each human centromere widely varies in length across different human genomes (Altemose et al. 2022) and it is just a coincidence that the first two assembled human centromeres have similar lengths.

### The key limitation of the standard sequence alignment

The orange path in Figure Alignment, left, shows the highest-scoring alignment path between centromeres *cenX*_*1*_ and *cenX*_*2*_ constructed by the standard dynamic programming algorithm for sequence comparison. This alignment suggests that these two centromeres are very similar with 98% percent identity (28,421 bp are insertions or deletions, 37,975 bp are mismatches, and 3,057,465 bp are matches). However, since centromeres widely vary in length across the human population (Altemose et al. 2022), this is likely a wrong conclusion that is affected by selecting two centromeres that coincidentally happened to be similar in length.

Below we demonstrate that the orange alignment path in Figure Alignment, left, does not reveal the true sequence of events on the evolutionary path between these centromeres. The blue path in Figure Alignment, left shows the rare-alignment path between *cenX*_*1*_ and *cenX*_*2*_ constructed by TandemAlignment. This path illustrates a very different and complex evolutionary scenario with only 1,954,622 matched positions and 2,335,879 insertions and deletions - surprisingly, the orange and blue paths in Figure Alignment, left, do not coincide at any of their matching positions!

**Figure Alignment.**
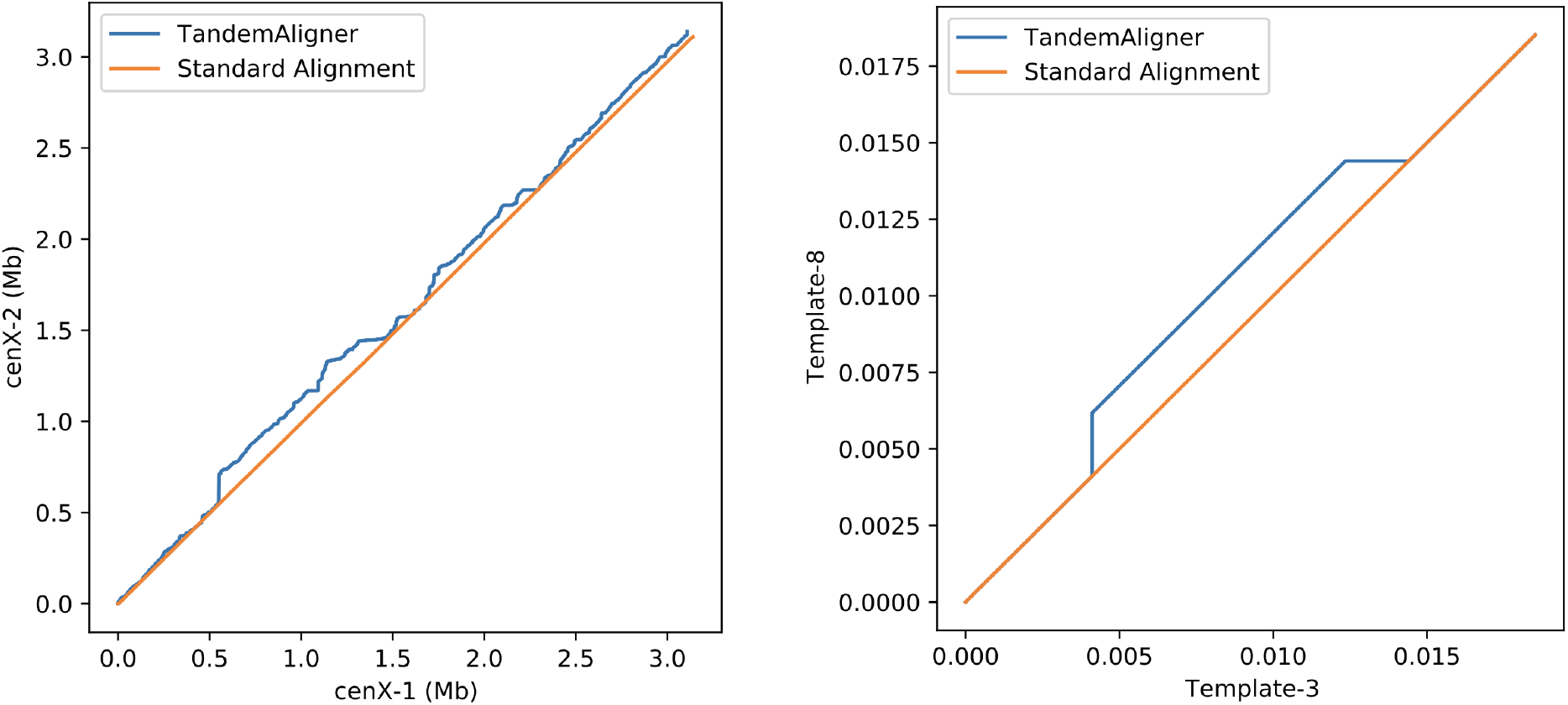
The highest-scoring alignment path between *c*entromeres *cenX*_*1*_ and *cenX*_*2*_ (left) and strings *Template*_*-3*_ and *Template*_*-8*_ (right) constructed by the standard alignment algorithm (orange path) and by TandemAligner (blue path). The blue path is not seen at positions where it coincides with the orange path. The standard alignment was constructed using the edlib tool (Šošić and Šikić 2017) with parameters match score 0, mismatch score −1, indel score −1 (changing parameters does not significantly change the alignment). The alignment of *Template*_*-3*_ and *Template*_*-8*_ is represented by the CIGAR string 4112M 1D 2354M 3I 2M 1I 7933M 1I 4111M. Other state-of-the-art sequence comparison tools result in similar alignments.

To illustrate limitations of the standard alignment using a simpler example, we generated the *HOR decomposition* of *cenX*_*1*_ (Kunyavskaya et al. 2022) and extracted ten HOR-blocks at coordinates *cenX*_*1*_:1,222,223-1,242,795 resulting in a sequence of length 20,572 bp referred to as a *Template*. We remove the third (eighth) HOR block at coordinates *Template:*4,112-6,172 (14,405-16,461), and refer to the resulting sequence as *Template*_*-3*_ (*Template*_*-8*_). The evolutionary correct alignment of *Template*_*-3*_ against *Template*_*-8*_ should align HORs 1, 2, 4, 5, 6, 7, 9, and 10, delete HOR 3, and insert HOR 8.

Sequences *Template*_*-3*_ and *Template*_*-8*_ are very similar (edit distance is 53 with only six gap-symbols) (Figure Alignment, right). Thus, the standard alignment fails to reveal the eight related HORs even in a simple case of *Template*_*-3*_ and *Template*_*-8*_ (let alone entire centromeres), illustrating that it does not adequately represent the correct evolutionary scenario in the case of repetitive sequences. Below we propose an alternative approach that leads to the correct alignment of *Template*_*-3*_ and *Template*_*-8*_. We first describe a simple scoring scheme that uses *rare k*-mers for a fixed *k-*mer size. TandemAligner improves on this scoring by using rare *k*-mers of varying sizes.

### Sequence-dependent alignment scoring based on rare *k*-mers

A *k*-mer in a string *S* is called *rare in S* if its count in *S* does not exceed a threshold *MaxCount*. A *k*-mer occurring in strings *S* and *T* is called *rare in S and T* if it is rare in both *S* and *T*. Other *k*-mers that are present in either *S* or *T* are referred to as *frequent*. Below we describe a simple alignment algorithm (referred to as TandemAligner_k_) based on sequence-dependent scoring that uses rare *k*-mers. TandemAligner is a parameter-free algorithm that improves TandemAligner_k_ by removing parameters *k* and *MaxCount*.

Given an alphabet *A*, we denote the alphabet of all *k*-mers from *A* as *A*^*k*^. Given a string *S* in the alphabet *A*, we denote the sequence of its |*S*|-*k*+1 *k*-mers (written in the alphabet *A*^*k*^) as *S*^*k*^. For example, for *S =* ACGT, *S*^*2*^=AC CG GT. TandemAligner_k_ transforms strings *S* and *T* into strings *S*^*k*^ and *T*^*k*^ and removes all frequent *k*-mers from these strings, resulting in strings *S*^*k\**^ and *T*^*k\**^ that consist of rare *k*-mers in *S* and *T*. Afterward, it constructs a *longest common subsequence* (*LCS*) between *S*^*k\**^ and *T*^*k\**^. Equivalently, TandemAligner_k_ finds an optimal alignment between *S*^*k*^ and *T*^*k*^ using the mismatch penalty equal to infinity and the indel penalty equal to zero. The premium for a match is *k*-mer-dependent and is defined as 1 (0) if the *k*-mer is rare (frequent) in *S* and *T*.

The described procedure constructs a *k*-mer-level alignment. Section “From *k*-mer-level alignment to regular nucleotide-level alignment” in Methods describes how to transform it into regular nucleotide-level alignment.

### Dot plots based on rare *k*-mers

Given a pair of sequences *S* and *T* and a string-set *X*, a *DotPlot*(*S, T*; *X*) is defined as a scatter plot such that each occurrence of every string *P* in *X* with starting coordinate *i* (*j*) in *S* (*T*) corresponds to a line connecting two points with coordinates (*i, j*) and (*i*+|*P*|, *j*+|*P*|). In the case when the string-set *X* consists of all rare *k*-mers in *S* and *T* for a threshold *MaxCount*, we refer to the set *DotPlot*(*S, T*; *X*) as *DotPlot*_*k,MaxCount*_(*S, T*). This definition of a dot plot differs from the standard definition as *DotPlot*_*k,MaxCount*_(*S, T*) only depicts rare *k*-mers shared by *S* and *T*, and reverse-complementary *k*-mers are treated as different *k*-mers.

Figure DotPlotsSynthetic present *DotPlot*_*k,MaxCount*_(*Template*_*-3*_, *Template*_*-8*_) for various values of parameters and illustrates that TandemAligner_k_ reveal the correct evolutionary scenario between *Template*_*-3*_ and *Template*_*-8*_ (except for the case *k*=10 and *MaxCount*=10 that we will address later). Figure DotPlotsCHM13HG002CenX shows dotplots *DotPlot*_*k,MaxCount*_(*cenX*_*1*_, *cenX*_*2*_) and the corresponding alignments constructed by TandemAligner_k_ (for various values of *k* and *MaxCount*) that reveal a complex evolutionary history of centromere X.

**Figure DotPlotsSynthetic.**
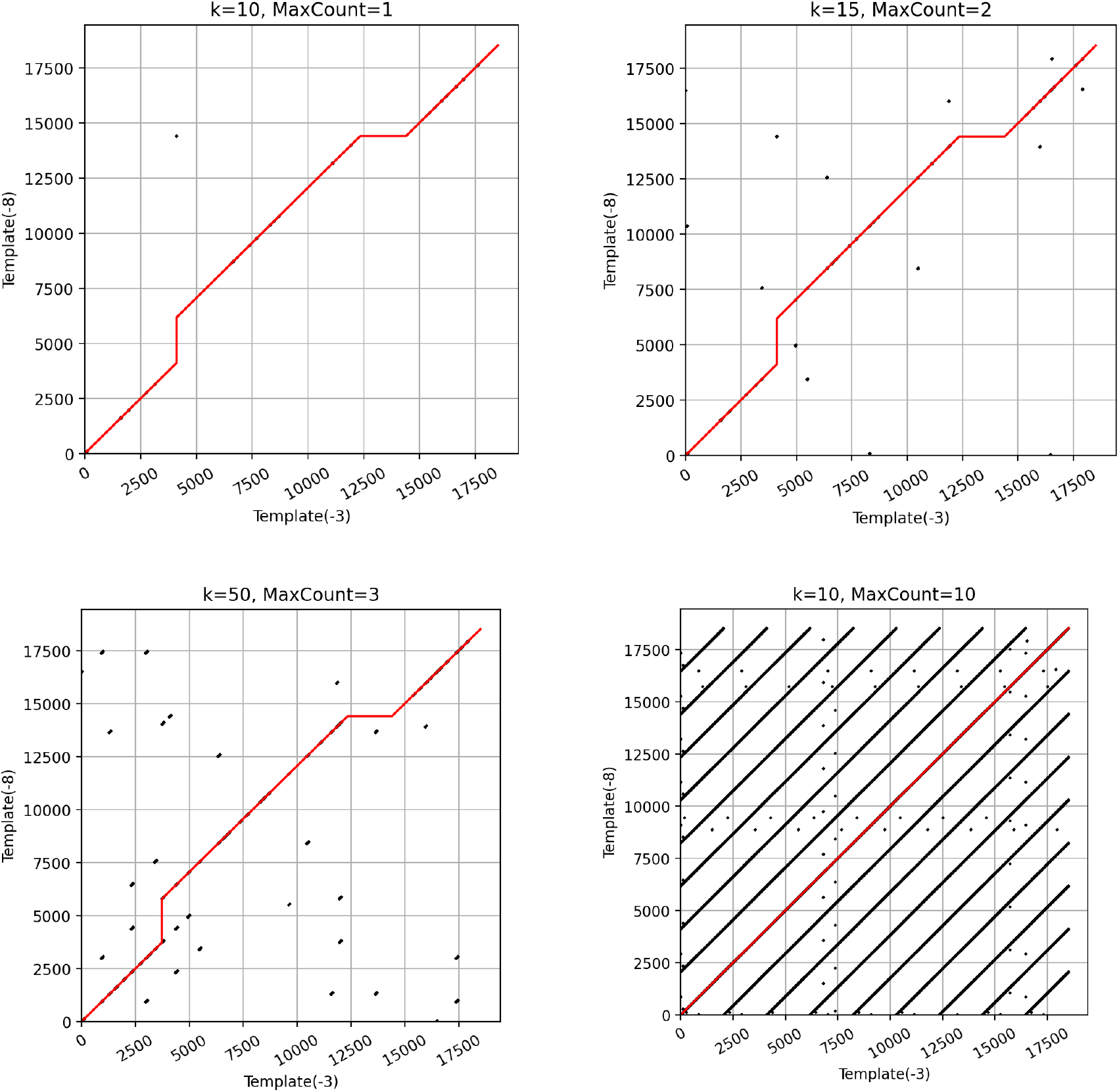
*DotPlot*_*k,MaxCount*_(*Template*_*-3*_, *Template*_*-8*_) for various values of parameters *k* and *MaxCount*. The red path corresponds to the optimal alignment constructed by TandemAligner_k_. In three cases, rare-alignment identifies the correct evolutionary scenario and reveals two indels that correspond to the third (eighth) HOR of *Template*_*-8*_.(*Template*_*-3*_) that is missing in *Template*_*-3*_ (*Template*_*-8*_). The parameters *k* = 10 and *MaxCount* = 10 result in an incorrect alignment that does not reveal the correct evolutionary scenario.

**Figure DotPlotsCHM13HG002CenX.**
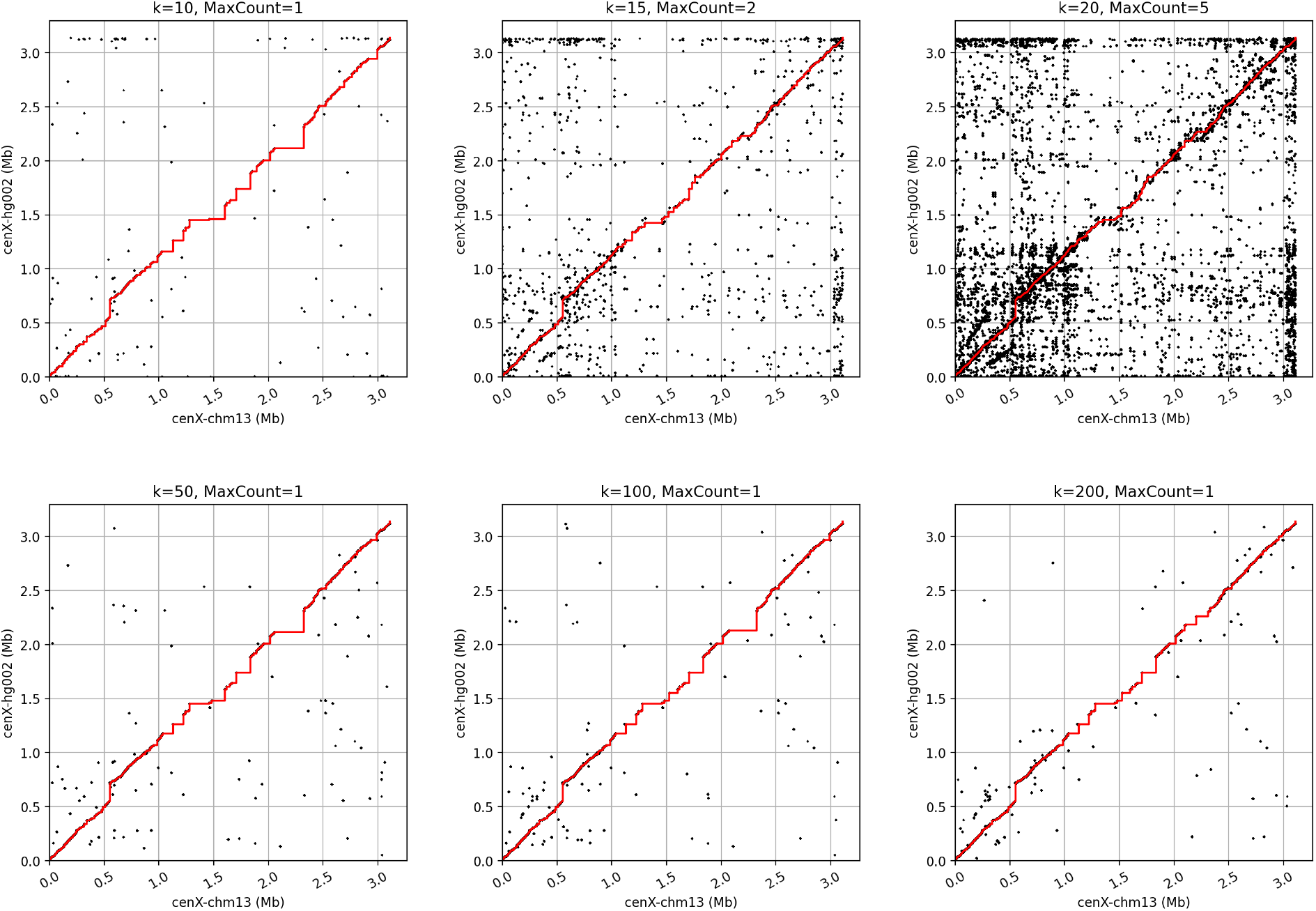
*DotPlot*_*k,MaxCount*_(*cenX*_*1*_, *cenX*_*2*_) for various values of parameters *k* and *MaxCount*. The red path corresponds to the optimal alignment constructed by TandemAligner_k_.

### Limitations of scoring based on rare *k*-mers of fixed size

Even though TandemAligner_k_ is a fast sequence alignment algorithm, it is unclear how to select a parameter *k*. On one hand, reducing its value ensures that one detects all shorter rare matches that cannot be extended either downstream or upstream. On the other hand, increasing the value of *k* provides reassurance that the detected matches are relevant to the evolutionary relationship between sequences rather than simply revealing a random coincident match. Simultaneously, a rare match that is longer than *k* might not contain any rare *k*-mers as its substrings and thus will not be detected. Even though, for a simple example in Figure DotPlotsSynthetic, such a dilemma is not a critical concern (except for the case *k*=10 and *MaxCount*=10), it remains unclear if an optimal choice for parameter *k* exists for real ETRs. Moreover, various regions of ETRs likely have varying optimal values of parameters *k*.

The last example in Figure DotPlotsSynthetic (*k*=10 and *MaxCount*=10) motivates development of a parameter-free alignment approach that does not rely on specific parameters *k* and *MaxCount*. TandemAligner improves on TandemAligner_k_ and reconstructs the correct evolutionary scenario between *Template*_*-3*_, and *Template*_*-8*_. by using an alternative parameter-free scoring approach.

### Shortest rare substrings

A substring *P* of a string *S* is called *n*-*rare* if *count*_*P*_(*S*) *= n* and *unique* if *count*_*P*_(*S*) = 1 (*count*_*P*_(*S*) refers to the number of occurrences of a substring *P* in a string *S*). For each *n* such that 1 ≤ *n* ≤ *MaxCount*, we consider the *shortest n*-rare substring starting at position *i* and denote its length as *rare*_*i*_(*S, n*). In the case when there is no *n*-rare substring starting at position *i*, we assign *rare*_*i*_(*S, n*)=∞ (there exists a 1-rare substring starting at each position since we add a unique “$” symbol to mark the end of a string). Figure RareLength illustrates that *rare*_*i*_(*cenX*_*1*_, 1) varies widely across the centromere. Positions located within recent duplications in cenX typically result in high values of *rare*_*i*_(*cenX*_*1*_, 1). Section “Algorithm for computing shortest rare substrings” in Methods describes an algorithm for computing *rare*_*i*_(*S, n*) for all 1 ≤ *i* ≤ |*S*| given a parameter *n* in O(|*S*|) time.

**Figure RareLength.**
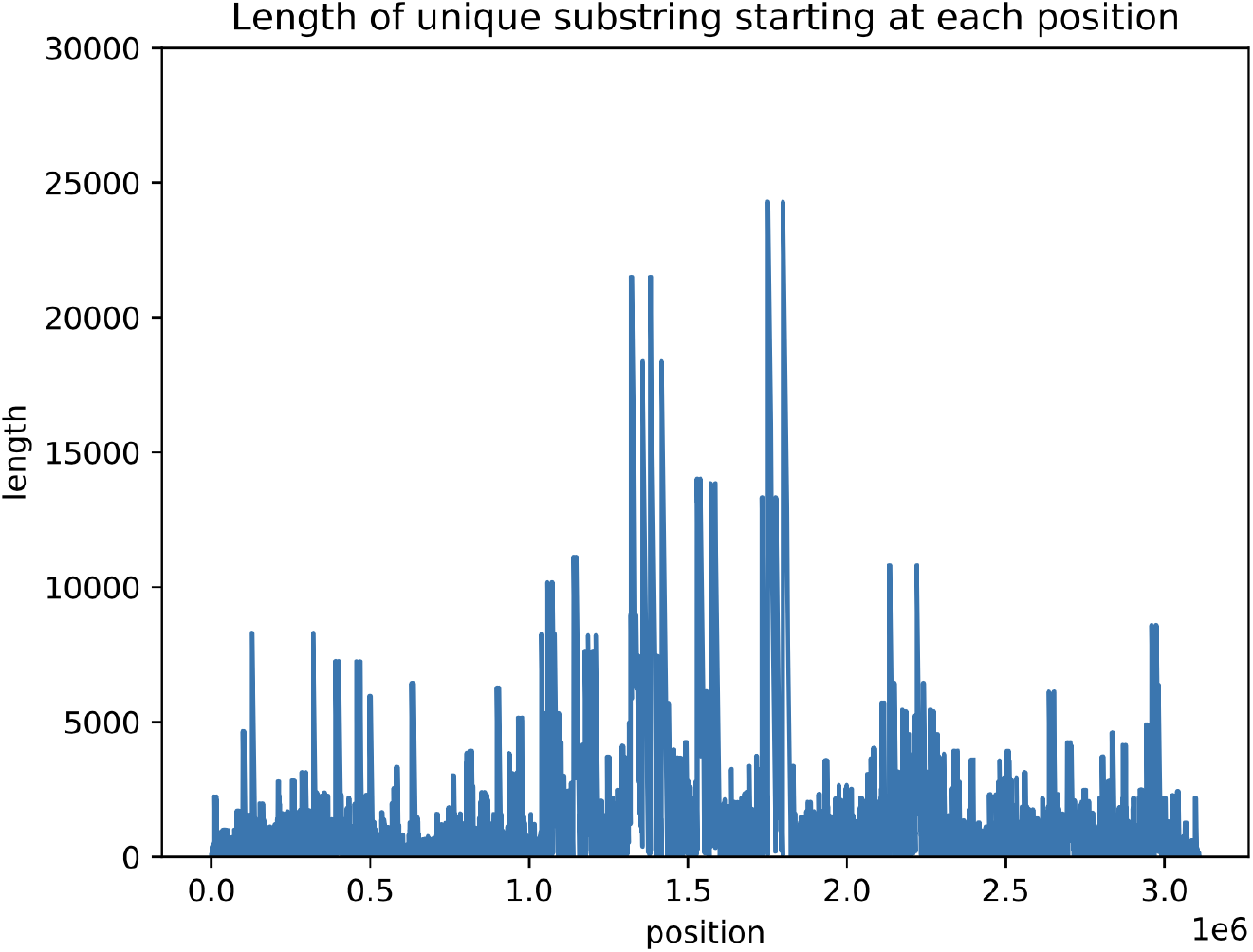
The array *rare*_*i*_(*cenX*_*1*_, 1). Although the mean value of *rare*_*i*_(*cenX*_*1*_, 1) is only 1,736, *rare*_*i*_(*cenX*_*1*_, 1) exceeds 10,000 for 94,976 positions in *cenX*_*1*_,

### Anchors

An *n*-rare substring *P* of *S* is called an *n-anchor* if no (proper) substring of *P* is an *n*-rare substring of *S*. Given a threshold *n* and array *rare*_*i*_(*S, n*) for all *i*, we can compute the set of the starting/ending positions *Anchor*_*n*_(*S*) of all *n*-anchors in O(|*S*|) time. Since the complexity of constructing the array *rare*_*i*_(*S, n*) is linear, the resulting complexity of finding the set *Anchor*_*n*_(*S*) in a string *S* is also O(|*S*|). Since the set of *n*-anchors is typically much smaller than the set of *n*-rare substring (e.g., cenX_1_ and cenX_2_ share only 9,708 1-anchors), TandemAligner uses an anchor-based scoring.

Given a string *S*, we use the shorthand *DotPlot*_*n*_(*S*) for *DotPlot*(*S, S, Anchors*_*n*_(*S*)). Figure DotPlotcenX combines the dotplots *DotPlot*_*n*_(*cenX*_*i*_) for *n*=2,3,4,5 and reveals the complex (and cryptic) evolutionary history of insertions/deletions in *cenX*_*1*_ and *cenX*_*2*_. Note that *DotPlot*_*n*_(*cenX*_*1*_) and *DotPlot*_*n*_(*cenX*_*2*_) share many anchors (reflecting duplications that happened before they diverged) but also a significant number of different anchors that reflect recent duplications in each of these centromeres.

**Figure DotPlotcenX.**
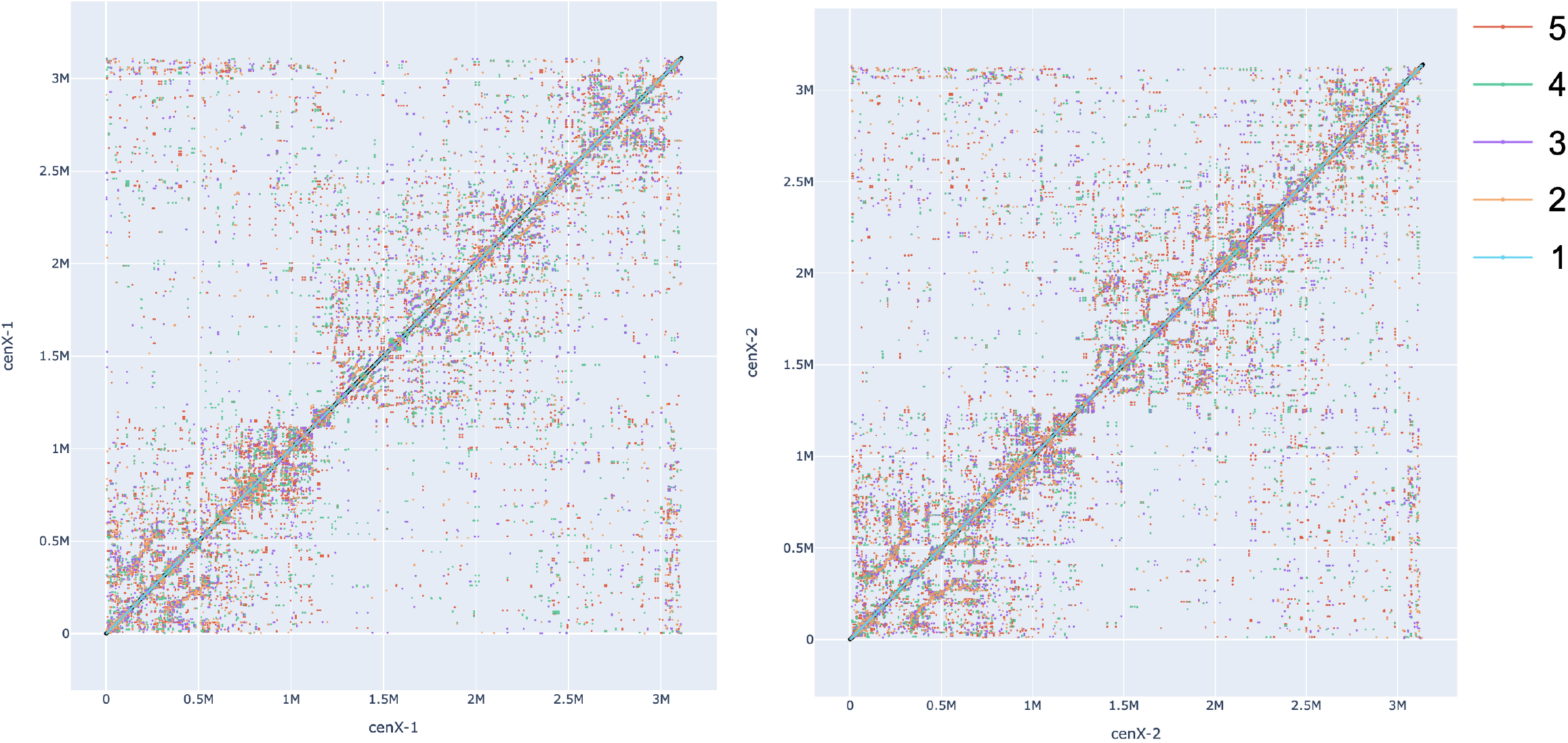
The superposition of *n*-rare dot plots *DotPlot*_*n*_(*cenX*_*1*_) (left) and *DotPlot*_*n*_(*cenX*_*2*_) (right) for *n*=2,3,4,5. Each of four dotplots is shown using an individual color.

Since two dot plots in Figure DotPlotcenX look similar (with clearly visible rectangles that have high density of points), one may arrive to a wrong conclusion that *cenX*_*1*_ and *cenX*_*2*_ have nearly identical architectures. Since the T2T Consortium did not have tools for deriving the detailed history of indels in centromere evolution it came to this conclusion by analyzing dot plots generated by the StainedGlass tool (Vollger, Kerpedjiev, et al. 2022). Specifically, it concluded that *cenX*_*1*_ and *cenX*_*2*_ are highly concordant apart from the three regions with recent insertions and deletions (Figure 5D in (Altemose et al. 2022). Below we show that *cenX*_*1*_ and *cenX*_*2*_ differ from each other by over 300 of large duplications and deletions of entire HORs (canonical HOR unit in centromere X has length 2057 bp). Moreover, six of them exceed 20 kb in length and include 11, 12, 20, 22, 25, and even 37 HOR units.

Given strings *S* and *T*, an (*n,m*)-*anchor* is an *n*-*anchor* in *S* and an *m*-*anchor* in *T*. We define *Anchors*_*n,m*_(*S,T*) *as* the set of all (*n,m*)-anchors in strings *S* and *T*. Finding the set *Anchors*_*n,m*_(*S,T*) for two strings *S* and *T* is analogous to finding the set of anchors for a single string and can be done in O(|*S*| + |*T*|) time. Figure DotPlotCen_1_-Cen_2_, left, combines *DotPlot*(*cenX*_*1*_,*cenX*_*2*_,*Anchors*_*n,m*_(*cenX*_*1*_,*cenX*_*2*_)) for four different values of *n* and *m*.

**Figure DotPlotCen1-Cen2.**
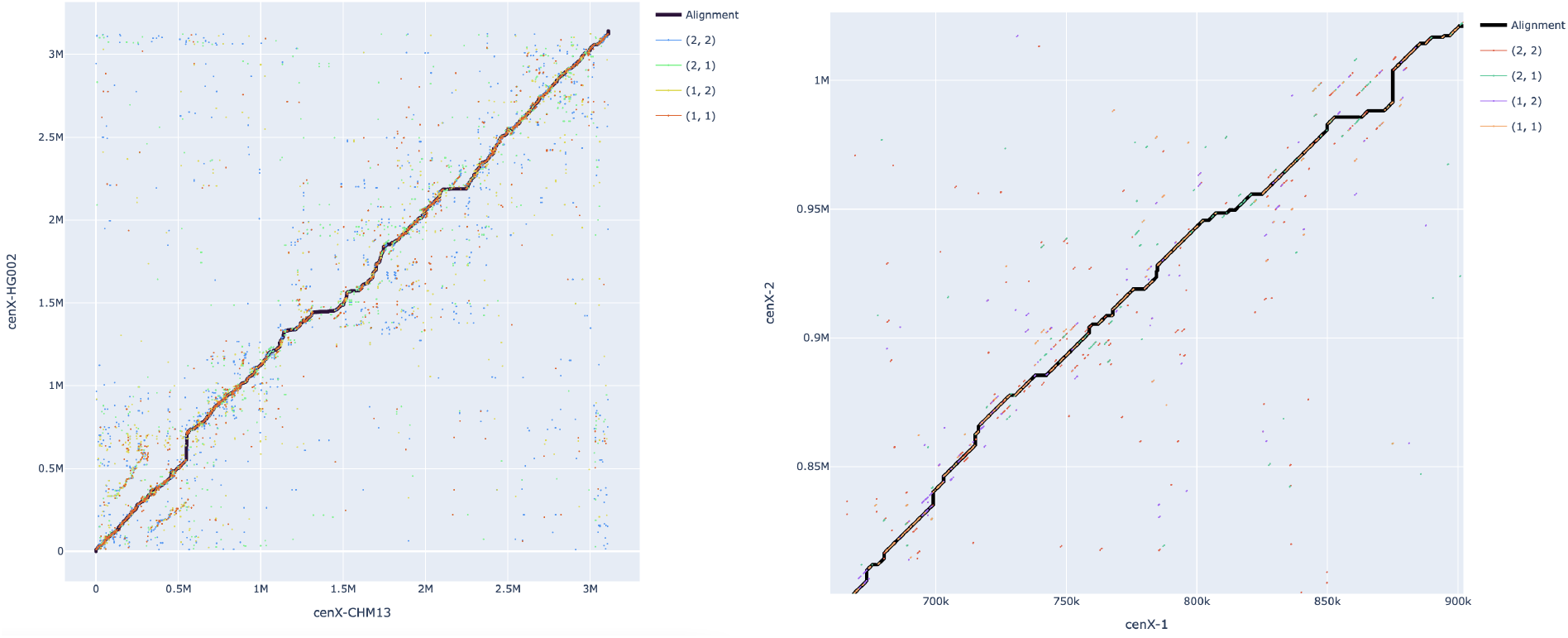
The anchor-based alignment of *cenX*_*1*_ and *cenX*_*2*_ and *DotPlot*(*cenX*_*1*_,*cenX*_*2*_,*Anchors*_*n,m*_(*cenX*_*1*_,*cenX*_*2*_)) for (*n,m*) = (1,1), (1,2), (2,1), (2,2). **(Left)** Anchor-based dot plot and alignment of the entire centromeres. **(Right)** Zoomed subrectangle shows the anchor-based dot plot of substrings of *cenX*_*1*_ and *cenX*_*2*_ of length approximately 220 kb and reveals that many lines in this dot plot aggregate into lines *j=i+t\**|*HOR*|.

### Anchor-based alignment graph

TandemAligner modifies the standard alignment graph (see section “Standard alignment graph” in Methods) by removing/adding some edges and modifying the scoring approach. First, we remove all diagonal edges from the graph and assign weight 0 to the remaining edges. Given a (positive) integer parameter *MaxCount*, we consider *AllAnchors = AllAnchors*(*S, T, MaxCount*) — the set of all occurrences of (*n, m*)-anchors in *S* and *T* for 1 ≤ *n, m* ≤ *MaxCount*. For an (*n,m*)-anchor *P*, such that *P* = *S*[*i*:*i*+|*P*|] = *T*[*j*:*j*+|*P*|], we add an edge connecting vertices (*i, j*) and (*i* + |*P*|, *j* + |*P*|) and assign it the weight equal to |*P*| / (*n*m*). We denote *K = ∑*_*P in AllAnchors*_ *count*_*P*_(*S*)\**count*_*P*_(*T*) *—* the number of diagonal edges in this graph.

The standard dynamic programming algorithm finds the heaviest path in this graph in O(|*S*|*|*T*|) time. However, since all vertical and horizontal edges have weight 0, one can find this path in O(*K*^*2*^) time by conducting the sparse dynamic programming computation only for the starting/ending vertex of each diagonal edge (Pearson and Lipman 1988). Although TandemAligner uses this simple approach, more advanced algorithms solve this problem in O(*K* * log *K*) time, resulting in further speed-up (Eppstein et al. 1992). In the case of centromere comparison, *K* is usually small, e.g., for rare-alignment of *cenX*_*1*_ against *cenX*_*2*_ with *MaxCount* = 1, *K* = 9708. The next subsection explains why such a low value of *K* is sufficient for constructing accurate alignments.

Since the set *AllAnchors* can be computed in O(*MaxCount*^2^ * (|*S*|+|*T*|)) time, the resulting running time of the anchor-based alignment algorithm with a single parameter *MaxCount* is O(*MaxCount*^2^ * (|*S*|+|*T*|) + *K*^2^). Figure DotPlotCen_1_-Cen_2_, left, shows an anchor-based alignment of *cenX*_*1*_ and *cenX*_*2*_.

### Recursive anchor-based rare-alignment

The only trade-off of the anchor-based alignment is the choice of parameter *MaxCount*. On one hand, a lower value of the parameter *MaxCount* leads to more *sparse* alignment with fewer matched bases. On another hand, higher values of the parameter *MaxCount* increase the number of (*n,m*)-anchors and introduce a computational burden since the number *K* of diagonal edges in the anchor-based alignment graph increases.

We call a sequence of consecutive matches (insertions, deletions) in a rare-alignment as a *match-run* (*insertion-run, deletion-run*). For example, the anchor-based alignment of *cenX*_*1*_ against *cenX*_*2*_ with *MaxCount* = 1(5), has 1,447,947 (1,790,434) matched bases and 865 (1223) match-runs, 1,692,054 (1,349,567) insertions and 811 (1141) insertion-runs, and 1,661,351 (1,318,864) deletions and 808 (1137) deletion-runs. The number of shared matching positions between these two alignments is 1,399,978.

To address the trade-off between small and large values of parameter *MaxCount*, TandemAligner constructs parameter-free *rare-alignment* by introducing a recursive strategy described in Methods. The rare-alignment of *cenX*_*1*_ against *cenX*_*2*_ has 1,983,431 (2744) matched bases (match-runs), 1,156,570 (2479) insertion bases (insertion-runs), and 1,125,867 (2467) deletion bases (deletion-runs). This rare-alignment accounts for 535,484 increase in the number of matched bases compared to the non-recursive anchor-based alignment (Figure RecursiveAlignment).

**Figure RecursiveAlignment.**
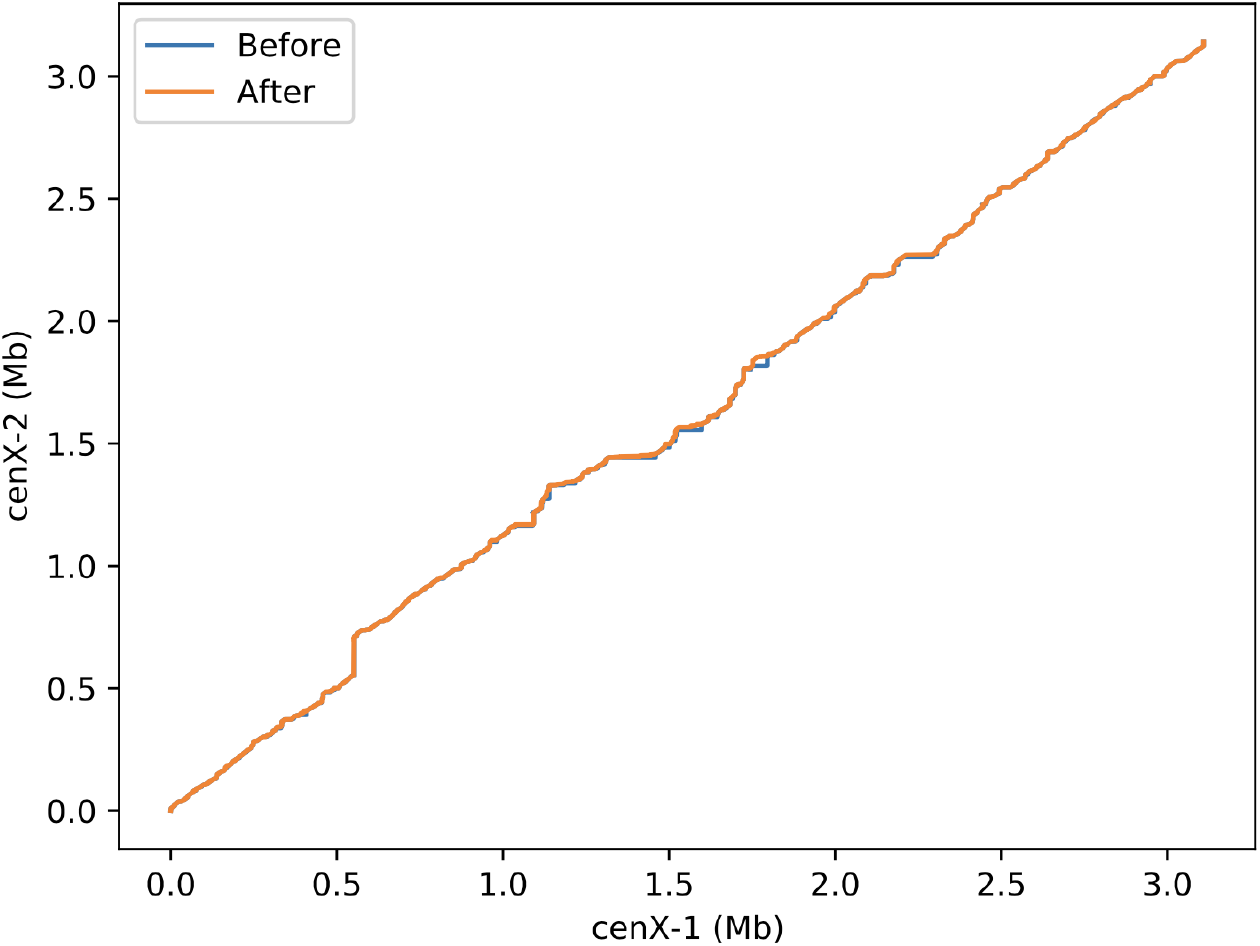
Anchor-based alignment path before applying the recursive procedure (blue) and rare-alignment after applying the recursive procedure (orange). The anchor-based alignment is constructed with parameter *MaxCount*=1. Although the changes appear to be small for a naked eye, the recursive procedure results in many changes that are difficult to see due to the small scale of the Figure.

The constructed recursive rare-alignment does not include any mismatches since it represents a single-nucleotide mismatch as a single-nucleotide insertion (deletion) edge followed by a single-nucleotide deletion (insertion) edge. Section “Refining rare-alignments” in Methods describes how TandemAligner transforms some of such pairs of indel edges into mismatches. Below we analyze the rare-alignment of centromeres and immunoglobulin loci. Further benchmarking is provided in Supplementary Notes “Benchmarking standard alignments and rare-alignments on simulated centromeres”, “Benchmarking standard alignments and rare-alignments on non-repetitive strings”.

### TandemAligner reveals the very high rate of large deletions and duplications in centromeres

Figure IndelHistogram, left shows the distribution of lengths of insertion-runs and deletion-runs in the rare-alignment of *cenX*_*1*_ and *cenX*_*2*_. Notably the prominent peaks in this distribution correspond to the length of a single canonical HOR in cenX (2057 bp) or to the lengths of multiple canonical HOR. The fact that TandemAligner automatically derived the HOR length in cenX without any prior knowledge adds confidence that it adequately represents evolution of centromeres.

For an indel *IND*, we compute *multiplicity*(*IND*) as the closest integer to |*IND*|/|*HOR*|, where |*IND*| is the length of the indel and |*HOR*| is the length of the canonical HOR (2057 bp for centromere X). We further define *offset*(*IND*)=||*IND*|-*multiplicity*(*IND*)*|*HOR*|| and classify an indel *IND* as a *HOR-indel* if *offset*(*IND*)/|*HOR*| does not exceed a threshold (the default value 0.05). The rare-alignment of *cenX*_*1*_ and *cenX*_*2*_ includes 175 (59, 28, 63) HOR-indels of multiplicity 1 (2, 3, more than 3) suggesting that HOR-indels formed by a single HOR dominate centromere evolution with 54% (18%, 9%, 19%) HOR-indels of multiplicity 1 (2, 3, more than 3). It will be interesting to see whether these numbers are stable for other centromeres across the human population when their sequences become available. Figure IndelHistogram, right shows distribution of HOR-indel over *cenX*_*1*_ and reveals their high density over the entire length of the centromere.

With about twenty thousands structural variations (SVs) per a single human genome, (Liao et al. 2022) the rate of SVs in the human genome is estimated as roughly one SV per 150 kb on average. With 325 HOR-indels identified by TandemAligner, the rate of SVs in human centromeres may be as large as one SV per 10 kb, an order of magnitude increase.

### TandemAligner reveals the low rate of single-nucleotide substitutions and small indels in centromeres

Although estimating the rate of single-nucleotide substitutions (and small indels) is a straightforward task for most genomic regions, it should be done with caution in ETRs. Constructing an accurate alignment of ETR (and limiting attention to regions of this alignment that do not include large indels) is a prerequisite for an accurate estimate of the single-nucleotide mutation rates in ETRs.

For example, the mutation rate between sequences *Template*_*-3*_ and *Template*_*-8*_ is zero since each of them was generated from the same sequence *Template* by a large deletion. However, the standard alignment (orange path in Figure Alignment, right) suggests that these sequences have 53 mutations, resulting in an inflated estimate of the mutation rate (53/18,500=0.0029) that exceeds the average mutation rate in the human genome. Similarly, the standard alignment of *cenX*_*1*_ against *cenX*_*2*_ (orange path in Figure Alignment, left) suggests that these sequences have an extremely high mutation rate (nearly 1% for single-nucleotide substitutions and over 1.2% for short indels) that exceeds the average mutation rate in the human genome by an order of magnitude. However, such high mutation rates (consistent with previous papers aimed at analyzing mutation rates in centromeres(Rudd et al. 2006; Pertile et al. 2009; Logsdon et al. 2021)) are merely artifacts of the incorrect alignments.

Moreover, even with a correct alignment, estimation of the mutation rates should be done with caution after limiting attention to the regions that do not include large indels. For example, after removing regions corresponding to two large indels in the rare-alignment between *Template*_*-3*_ and *Template*_*-8*_ (blue path in Figure Alignment, right), we are left with identical regions that result in the correct 0% estimate of the mutation rate between *Template*_*-3*_ and *Template*_*-8*_.

We call an indel *short* if its length does not exceed a threshold *ShortIndel*, and *long*, otherwise (default value *ShortIndel =* 5). The rare-alignment of *cenX*_*1*_ and *cenX*_*2*_ contains 268 short indels (of total length 473) and 520 long indels (of total length 2,277,742). Removing all long indels from the alignment-path of *cenX*_*1*_ and *cenX*_*2*_ breaks it into a set of shorter paths (with total length *L*^*-*^ =1,986,015) that contains *M*=2107 mismatches and 268 short indels of total length *I*=473. We use this path-set to estimate the rate of mismatches and (short) indels as *M*/ *L*^*-*^ ≅1 / 1000 bp and *I/L*^*-*^≅2 / 10,000, respectively. Thus, analysis of rare-alignments suggests that previous studies likely came to the incorrect conclusion that the rate of single-nucleotide mutations is greatly elevated in centromeres (Rudd et al. 2006; Pertile et al. 2009; Logsdon et al. 2021).

**Figure IndelHistogram.**
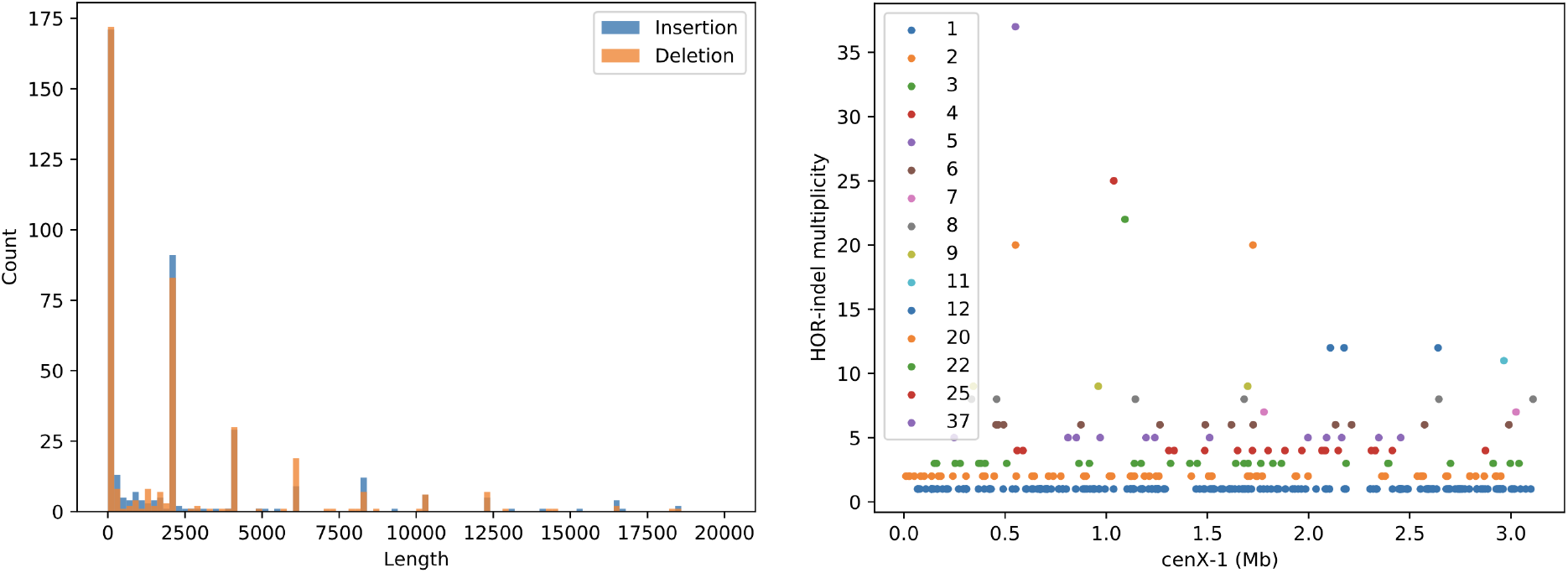
The histogram of lengths of insertion-runs and deletion-runs in the rare-alignment of *cenX*_*1*_ and *cenX*_*2*_ (left) and distribution of HOR-indels in this alignment along the entire length of *cenX*_*1*_. **(Right)** The peaks in the histogram correspond to either short (< 100bp) runs or to likely insertions / deletions of a single or multiple canonical HOR units of cenX. The length of the canonical HOR in cenX is 2057 bp (Miga et al. 2020), while the peaks in the histogram correspond to 2057 bp (a single HOR unit), 4114 bp (two HOR units), 6171 bp (three HOR units), 8225 bp (approximately 4 HOR units), etc. The width of each bar is 500bp. Even though the histogram’s *x*-axis is cut at 20 kb, there are long indels of lengths 22,628 bp (11 HOR units), 24,676 bp (12 HOR units), 41,133 bp (20 HOR units), 45,225 (22 HOR units), 51,393 bp (25 HOR units), and 76,098 bp (37 HOR units). (**Left**) Each color corresponds to an indel of a particular length, e.g., blue (orange, green) color corresponds to HOR-indel of multiplicity 1 (2, 3), etc.

### TandemAligner reveals orthologous D genes in primate immunoglobulin loci

Comparative analysis of the immunoglobulin genes across multiple species is a prerequisite for evolutionary studies of the adaptive immune system in vertebrates. Below we focus on D genes in the immunoglobulin IGHD locus that play a key role in diversifying antibody repertoires and that are notoriously difficult to compare between species and predict in newly sequenced genomes (Sirupurapu, Safonova, and Pevzner 2022). Moreover, since D genes are located within ETRs, identifying pairs of orthologous D genes even between close species (such as primates) is challenging.

This difficulty is further compounded by the fact that assembly of the highly-repetitive immunoglobulin loci has been challenging (Watson and Breden 2012; Rodriguez et al. 2020). However, even though there were very few genomes available for comparative immunogenomics studies until recently, the situation has changed with the advent of contiguous long-read assemblies generated by the Vertebrate Genomes Project (VGP) (Rhie et al. 2021). Nevertheless, the existing methods for finding orthologous genes (Koonin 2003) are not well-suited for D genes since these methods are typically based on similarity search that often fails to detect similarities between highly diverged and short D genes (most are shorter than 50 bp) located within complex ETRs. As a result, the IGHD loci in hundreds vertebrate genomes recently assembled by the VGP consortium remain poorly annotated (Sirupurapu, Safonova, and Pevzner 2022).

The IGHD locus in mammalian genomes includes a long tandem repeat (Safonova and Pevzner 2020). For example, the human IGHD locus contains a tandem repeat formed by four ∼10 kb long units while the orangutan IGHD locus contains a tandem repeat formed by five ∼10 kb long units. Finding orthologs between these units (and thus analyzing evolution of D genes) is challenging,

Below we benchmark TandemAligner and edlib on human and orangutan IGHD loci. We extracted the region of the heavy immunoglobulin locus of the human genome (referred to as *IGHD*_*H*_; NC_000014.9:105,865,737-105,964,717) and orangutan genome (referred to as *IGHD*_*O*_; NC_036917.1:87,805,787-87,899,312) that contains D genes. Both regions contain long tandem duplications that differ in the number of units (four for human and five for orangutan). Thus, the evolutionary correct alignment should match the first *i* units, followed by a unit-insertion in orangutan genome (or a unit deletion in human genome), followed by matches of the remaining 4-*i* units (*i* is unknown). In difference from edlib, TandemAligner reveals the additional unit of the tandem repeat in the orangutan IGHD locus and suggests that this additional unit emerged in the end of the IGHD locus (*i*=4) in the common ancestor of humans and orangutans, thus establishing orthology between human and orangutan D genes (Figure IGHDLocus).

**Figure IGHDLocus.**
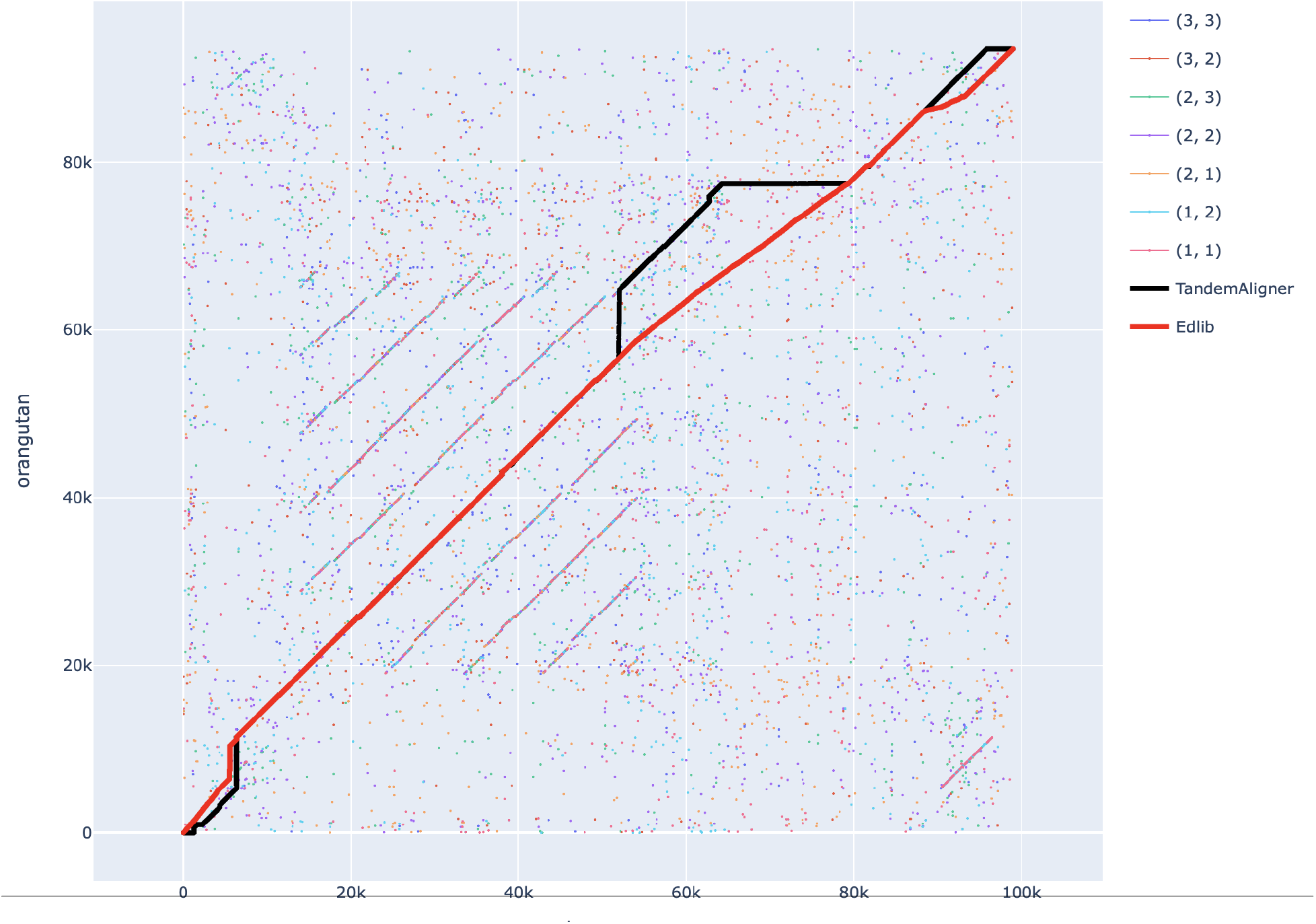
**The rare-alignment (black) and standard alignment (red) of strings *IGHD***_***H***_ **and *IGHD***_***O***_ **and *DotPlot*(*IGHD***_***H***_,***IGHD***_***O***_,***Anchors***_***n***,***m***_**(*IGHD***_***H***_,***IGHD***_***O***_**)) for (*n***,***m*) = (1**,**1), (1**,**2), (2**,**1), (2**,**2), (2, 3), (3, 2), (3, 3)**.

### Running time of TandemAligner

It takes TandemAligner 25 seconds to compute a rare-alignment and generate the CIGAR-string for *cenX*_*1*_ and *cenX*_*2*_. For comparison, it takes Edlib — a state-of-the-art fast alignment algorithm (Šošić and Šikić 2017) — 58 seconds on the same machine. Both tools use a single CPU thread.

## Methods

### From *k*-mer-level alignment to regular nucleotide-level alignment

A *k*-mer-level alignment between strings *S*^*k*^ and *T*^*k*^ induces a regular nucleotide-level alignment. Each rare *k*-mer in the alignment of strings *S*^*k*^ and *T*^*k*^ induces *k* matches (*i,j*), (*i+*1,*j+*1), …,(*i+k-*1,*j+k-*1) between individual positions in strings *S* and *T*. Since matches of overlapping *k*-mers may induce multiple matches of a single position in the string *S* (*T*), we need to remove some of the induced matches to ensure that a single position in one string is matched to a single position in another string. We say that an induced match (*i,j*) *precedes* induced matches (*i, j+Δ*) and (*i+Δ, j*) if *Δ*>0 and iteratively remove all matches that are preceded by other matches. The remaining matches (along with resulting unmatched symbols that form indels) form a nucleotide-level alignment between strings *S* and *T*.

### Algorithm for computing shortest rare substrings

Below we describe an algorithm for computing the array *rare*_*i*_(*S, n*) for all 1 ≤ *i* ≤ |*S*| given a parameter *n* in O(|*S*|) time.

Let *S*(*i*) be the suffix corresponding to the *i*-th element of the suffix array of the string *S* (Manber and Myers 1989). We define *LCP*(*k, l*) as the Longest Common Prefix of *S*(*k*) and *S*(*l*)). TandemAligner first computes the standard Longest Common Prefix (LCP) array *LCP*(*i*-1, *i*) for consecutive elements *i-*1 and *i* in the suffix array and uses it to construct the array LCP(*k,k+n*-1). It computes each element of this array in O(1) time by iterating through the standard LCP array and using a deque with a minimum over the standard LCP array.

A segment [*k, l*] is called *n-rare* if LCP(*k,l*) is larger than both LCP(*k*-1, *l*) and LCP(*k, l*+1) and *l-k*+1 = *n*. Given an index *i*, we find the *n*-rare segment [*k, k+n*-1] containing *S*(*i*). Finally, we set *rare*_*i*_(*S, n*) = 1 + max(LCP(*k*-1, *l*), LCP(*k, l*+1)). If no such rare segment exists, suffix *S*(*i*) itself is frequent. In this case, we set *rare*_*i*_(*S, n*) = ∞. Since we can compute *rare*_*i*_(*S, n*) for all *i* during a single iteration through the LCP(*k,k+n*-1) array (and the elements of this array can be efficiently computed from the standard LCP array), the complexity of calculating *rare*_*i*_(*S, n*) for all *i* is O(|*S*|). Since the LCP array can be constructed from the suffix array in O(|*S*|) time (Kasai et al. 2001), the complexity of building *rare*_*i*_(*S, n*) for a string *S* is O(|*S*|). In practice, we use a fast suffix array construction algorithm with complexity O(|*S*| * log |*S*|) (Jesper Larsson 1999; Burkhardt and Kärkkäinen 2003) rather than O(|*S*|) which is sufficient for practical purposes (Kärkkäinen and Sanders 2003; Kim et al. 2003; Ko and Aluru 2003).

### Standard alignment graph

The standard weighted directed acyclic graph that is used for the alignment of strings *S* and *T* presents a grid with |*S*|+1 rows and |*T*|+1 columns. For 1 ≤ *i* ≤ |*S*| and 1 ≤ *j* ≤ |*T*|, the vertex with coordinate (*i, j*) is connected via a *vertical, diagonal*, and a *horizontal* edge to vertices (*i*+1, *j*), (*i*+1, *j*+1), (*i, j*+1), respectively. Vertex (|*S*| + 1, *j*) in the last “row” of the grid is connected to the vertex (|*S*| + 1, *j* + 1) via a horizontal edge for 1 ≤ *j* ≤ |*T*|, and vertex (*i*, |*T*|+1) in the last “column” of the grid is connected to (*i* + 1, |*T*| + 1) via a vertical edge for 1 ≤ *i* ≤ |*S*|. The diagonal edge connecting vertices (*i, j*) and (*i*+1, *j*+1) corresponds to a *match* of characters *S*[*i*] and *T*[*j*] in case they are equal or to a *mismatch*, otherwise. The vertical edge connecting vertices (*i, j*) and (*i*+1, *j*) corresponds to an *insertion* of character *S*[*i*] between *T*[*j*] and *T*[*j*+1]. Horizontal edge connecting vertices (*i, j*) and (*i, j*+1) corresponds to a *deletion* of the character *T*[*j*] between *S*[*i*] and *S*[*i*+1]. The simplest scoring strategy in the standard alignment approach assigns the match (mismatch) weight to all diagonal edges that define matches (mismatches), and indel weight to all vertical and horizontal edges.

### Recursive algorithm for constructing rare-alignment

Given strings *S* and *T*, TandemAligner constructs the anchor-based alignment *Alignment* using the set *Anchors*(*S, T*, 1) or, if this set is empty, for the minimal value of *Count* such that this set is non-empty. We refer to a segment of *Alignment* consisting of a deletion-run immediately followed by an insertion-run (or vice versa) as an *indel-pair*. Each indel-pair corresponds to a pair of substrings *s* and *t* in *S* and *T*, respectively. TandemAligner recursively constructs an alignment on strings *s* and *t*, and substitutes the indel-pair in *Alignment* by the resulting (small) alignment-path. The process terminates when there are either no indel-pairs left or the set of anchors constructed on substrings corresponding to each remaining indel-pair is empty for any choice of *Count*. In practice, we limit the value of *Count* by a large constant *MaxCount* (default value = 50) since matches of substrings of higher counts are likely to be spurious. However, this parameter is simply a practical heuristic that is not necessary for the theoretical description of the algorithm.

### Refining rare-alignments

The recursive rare-alignment does not include any mismatches since it represents a single-nucleotide mismatch as an insertion (deletion) edge followed by a deletion (insertion) edge. Given a rare-alignment *Alignment*, TandemAligner searches for *square* indel-pairs that contain an *equal* number of insertions and deletions and represent likely runs of mismatches. For a given square indel-pair, we refer to this number as the *length* of this block. Recursive anchor-based rare-alignment of *cenX*_*1*_ and *cenX*_*2*_ contains 2051 (24, 4) square indel-pairs of length 1 (2, 3). For each square indel-pair in *Alignment*, TandemAligner substitutes it by a diagonal series of matches/mismatches in *Alignment*. Such a procedure applied to the recursive anchor-based rare-alignment of *cenX*_*1*_ and *cenX*_*2*_ introduces only 4 additional matches. The number of insertions (insertion-runs) reduced by 2111 (2079) to 1,154,459 (400). The number of deletions (deletion-runs) reduced (by the same value) to 1,123,756 (388). The number of introduced mismatches (mismatch-runs) is 2107 (2083).

## Discussion

The ongoing effort to construct the human pangenome promises to change the way we analyze genomic variations and infer their associations with diseases. The construction of the pangenome graph is based on generating alignments between complete human genomes that are now being assembled by the HPR Consortium (Wang et al. 2022). Although these alignments have already been constructed for the vast majority of regions of the human genome, (Liao et al. 2022) it turned out that the standard alignment approach fails to adequately align the most repetitive (and biomedically important) regions such as centromeres. The emerging consensus is that aligning such regions requires a new framework for sequence comparison rather than simply tweaking parameters of the standard alignment approach. As stated in (Liao et al. 2022): “*Our near-term goal is … to refine the pangenome alignment methods (so that telomere-to-telomere alignment is possible capturing more complex regions of the genome)*.”

We described a new parameter-free framework for sequence comparison and demonstrated that it reveals the evolutionary history of highly-repetitive regions such as centromeres and immunoglobulin loci. Admittedly, since only two human centromeres have been assembled, carefully validated, and publicly released so far, our benchmarking remains limited to these centromeres and various simulated examples. However, even this limited benchmarking shed light on the evolution of human centromeres (and the extremely high rate of duplications/deletions of single/multiple HORs) that the standard alignment approach failed to uncover.

Although TandemAligner is already fast, we plan to further speed it up using the fast sparse dynamic programming algorithm by Eppstein et al., 1992. Although the rare-alignment framework represents the first step toward aligning centromeres and other ETRs, many questions about human ETRs remain unanswered. For example, TandemAligner reveals extremely high-rate of large insertion-runs in human centromeres but does not answer the question which region of the ancestral centromeres contributed to each of these insertion-runs. Zooming on Figure DotPlotCen_1_-Cen_2_, right reveals that many lines in this dot-plot aggregate into lines *j=i+t\**|*HOR*| that are parallel to the main diagonal (|*HOR*| refers to the length of HOR in cenX and *t* is an integer). Since such aggregates are often triggered by HOR-insertions, their analysis should provide the initial clues for the common patterns of duplications in centromeres. Our next goal is to integrate analysis of duplications in the TandemAligner framework. Another bottleneck in applications of rare-alignments is that, in contrast to the analysis of statistical significance of standard alignments (Arratia and Waterman 1994; Waterman and Vingron 1994), it remains unclear how to estimate the statistical significance of rare-alignments (see Supplementary Notes “Summary of differences between standard alignment and rare-alignment”, “Evolution and alignment from probabilistic perspective”).

## Acknowledgements

We are grateful to Ivan Alexadrov, Anton Bankevich, Alexander Bzikadze, Tatiana Dvorkina, Olga Kunyavskaya, and Cynthia Wu for helpful discussions and suggestions.

## Authors’ contributions

A.V.B. conducted the experiments and wrote the code for TandemAligner. P.A.P. supervised the research. All authors worked on the development of the TandemAligner algorithm, wrote and edited the manuscript.

## Data Availability

Alignment of cenX_1_ and cenX_2_ generated by TandemAligner is located at Zenodo: https://zenodo.org/record/7058133.

## Supplemental Notes

- Benchmarking standard alignments and rare-alignments on simulated centromeres
- Benchmarking standard alignments and rare-alignments on non-repetitive strings
- Summary of differences between standard alignment and rare-alignment
- Evolution and alignment from probabilistic perspective

## Supplemental Note: Benchmarking standard alignments and rare-alignments on simulated centromeres

In addition to aligning *cenX*_*1*_ against *cenX*_*2*_, we aligned *cenX*_*1*_ against multiple centromeres simulated from *cenX*_*1*_ using both single-nucleotide mutations/indels and HOR-indels at various rates.

We quantify similarity between alignments as follows. Given an alignment-path *P*, we define *match*(*P*) as the set of all positions (*i,j*) on this path that represent matches. Given two alignment-paths *P* and *P’* of the same strings, we define *match*(*P,P’*) as the overlap between the set *match*(*P*) and *match*(*P’*). The similarity *sim*(*P,P’*) between alignments *P* and *P’* is defined as |*match*(*P,P’*)|/min{|*match*(*P*)|, |*match*(*P’*)|}.

To generate realistic simulated centromeres, we partitioned *cenX*_*1*_ into HORs (Kunyavskaya et al. 2022) and introduced *N* HOR-indel-runs. For the length of introduced HOR-indels, we use 1-based Poisson distribution with the mean estimated from Figure IndelHistogram, left and equal to 1.66. That results in 52%, 34%, 11%, 2.5%, and 0.5% of these HOR-indels representing HOR-indels of multiplicity 1, 2, 3, 4, and at least 5. These simulations (with the same rate of HOR-insertions and HOR-deletions) resulted in five simulated centromeres for *N*=100, 200, 500, 1000, 1500 with progressively increasing divergence from *cenX*_*1*_. To analyze centromeres of widely different sizes (that are common in human population), we also simulated five centromeres for the same values of *N* but this time with different rates of HOR-insertions and HOR-deletions (80% of HOR-insertions and 20% of HOR-deletions). We further simulated single-nucleotide mismatches and single-nucleotide indels (with estimated rates 0.1% and 0.02%, respectively) in each of the ten simulated centromeres, resulting in centromeres *c*_*100*,_ *c*_*200*,_ *c*_*500*,_ *c*_*1000*,_ *c*_*1500*_ and *c\**_*100*,_ *c\**_*200*,_ *c\**_*500*,_ *c\**_*1000*,_ *c\**_*1500*_.

Table CentromereBenchmarking illustrates that, in contrast to standard alignments, the rare-alignments constructed by TandemAligner approximate the correct alignment well.

**Table CentromereBenchmarking.** Similarity between correct alignments and alignments constructed by TandemAligner and edlib for ten datasets simulated from *cenX*_*1*_. Column “Length” corresponds to the length of simulated centromeres *c*_*100*,_ *c*_*200*,_ *c*_*500*,_ *c*_*1000*,_ *c*_*1500*_ and *c\**_*100*,_ *c\**_*200*,_ *c\**_*500*,_ *c\**_*1000*,_ *c\**_*1500*_. TandemAligner improves on edlib in all cases.

| $N$ | 50% of HOR-insertions and HOR-deletions | | | 80% of HOR-insertions and 20% HOR-deletions | | |
| --- | --- | --- | --- | --- | --- | --- |
|  | Length | TandemAligner | Edlib | Length | TandemAligner | Edlib |
| 100 | 3,120,950 | 0.96 | 0.28 | 3,303,647 | 0.96 | 0.85 |
| 200 | 3,113,239 | 0.93 | 0.14 | 3,557,802 | 0.92 | 0.66 |
| 500 | 3,110,073 | 0.84 | 0.06 | 4,114,758 | 0.81 | 0.63 |
| 1000 | 3,157,335 | 0.76 | 0.05 | 5,065,103 | 0.69 | 0.49 |
| 1500 | 3,106,405 | 0.69 | 0.03 | 6,125,774 | 0.64 | 0.33 |

## Supplemental Note: Benchmarking standard alignments and rare-alignments on non-repetitive strings

Similarly to benchmarking conducted by the authors of the state-of-the-art fast alignment tools SeqAn (Döring et al. 2008) and edlib (Šošić and Šikić 2017), we analyzed TandemAligner’s performance on non-repetitive real and simulated nucleotide sequences. First, we launched TandemAligner and edlib to compare genomes of two human influenza viruses *Virus* and *Virus’* (RefSeq GCF_001343785.1 and GCF_000865725.1, each of length approximately 13 kbp). The similarity between alignments constructed by TandemAligner and edlib is 87%.

Afterward, we simulated mismatches and single-nucleotide indels in *Virus* with the same mismatch and indel rates such that the combined mutation rate *MutRate* is equal to 2%, 5%, 10%, 15%, 20%, 25%, and 30%. We further introduced a deletion of length 1 kb at a randomly chosen position in each of these sequences, resulting in seven simulated sequences *Virus*_*mutRate*_.

Table edlibComparison presents benchmarking results for TandemAligner and edlib. We computed the percentage of correctly matched bases between *Virus* and *Virus*_*mutRate*_ in alignments constructed by TandemAligner and edlib using the concept of similarity between alignments defined in Supplemental Note “Benchmarking standard and rare-alignments on simulated centromeres”.

**Table edlibComparison.** Benchmarking TandemAligner and edlib on non-repetitive sequences. Similarity (range 0 to 1) between the ground truth alignment of *Virus* and *Virus*_*mutRate*_ and the alignment reported by each algorithm. Each row corresponds to a particular *Virus*_*mutRate*_. The best value for each row is highlighted in bold.

| Mutation Rate | Similarity between alignment |  |
| --- | --- | --- |
|  | TandemAligner | Edlib |
| 2% | **0.997** | 0.97 |
| 5% | **0.99** | 0.96 |
| 10% | **0.96** | 0.94 |
| 15% | 0.92 | **0.93** |
| 20% | 0.86 | **0.90** |
| 25% | 0.82 | **0.88** |
| 30% | 0.74 | **0.85** |

## Supplemental Note: Summary of differences between standard alignment and rare-alignment

To illustrate the differences between standard alignment and rare-alignment, we consider strings over a large alphabet *A* and assume that various symbols in these strings have widely different counts (similarly to different *k*-mers in a centromere *C* that have widely different counts in a string *C*^k^). Each alignment of strings *S* and *T* in this alphabet corresponds to a path in the standard alignment graph with (|*S*|+1)*(|*T*|+1) vertices with diagonal (matches and mismatches), horizontal (insertions), and vertical (deletions) edges. We assign labels to all edges as follows: each match edge is labeled by a symbol that represents the corresponding matching symbols in *S* and *T* and all other edges are labeled by a special symbol “#”. The score of an edge (interpreted as its “probability”) is defined as a function *p*(*a*) on symbols from the alphabet *A* (0 <= *p*(*a*) <=1) and as *p*(#)=1. Note that, for a given vertex, “probabilities” of all outgoing edges do not sum up to a unit, but can be appropriately normalized. The score of a path (alignment) is defined as the product of scores of its edges. This is a highly simplified framework for describing the standard alignment generalizing the longest common subsequence problem (by varying scores of different matching symbols) that ignores mismatch and indel penalty but captures the essence of this problem.

Both standard alignment and rare-alignment translate into a problem of finding a minimum-score path in the alignment graph but differ in the way the scoring function *p*(*a*) is defined. In bioinformatics applications, the choice of this function is critically important for the resulting alignment being biologically adequate. The key limitation of standard alignment (at least with respect to aligning ETRs) is that it assigns scoring parameters *p*(*a*) even before “seeing” the sequences *S* and *T*. Another limitation is that it is unclear how to select these scoring parameters that will work across all sequences.

The rare-alignment represents a more adequate probabilistic model since it instead defines *p*(*a*) as the probability that symbol *a* appears at two randomly chosen positions in *S* and *T*, i.e., as (*count*_*a*_(*S*)\**count*_*a*_(*T*))/|*S*|*(|*T*|). Given an alignment path with match-edges labeled by symbols *a*_1_,…,*a*_n_, we can assume that observing labels *a*_1_,…,*a*_n_ on these edges equals to the products *p*(*a*_1_)*…\**p*(*a*_n_) if we make a simplifying (and incorrect) assumption that these events are independent. An accurate way to compute this probability is to consider an urn model with balls of |*A*|+1 colors (one of them corresponds to the symbol “#”) but it is unclear how to effectively incorporate this more accurate scoring into the dynamic programming algorithm on the alignment graph. However, even though the score of the rare-alignment represents an approximation in this simplified model, it may not significantly affect relative ranking of various high-scoring paths and, as we demonstrated above, generates adequate alignments, at least for simulated examples.

## Supplementary Note: Evolution and alignment from probabilistic perspective

### Alignment as a trajectory of a Markov Chain

We will now show the connection between dynamic programming and Markov Chains. Let *G* be a weighted directed acyclic graph with edge-weights being arbitrary real numbers. For a vertex *v*, we transform edge-weights *w*_*1*_, *w*_*2*_, …, *w*_*v*_ for all outgoing edges of *v* as follows. We apply a *soft-max transformation (Bridle 1990)* to these weights to ensure that the transformed values are non-negative and sum up to a unit, thus, defining a probability distribution. Formally, weight *w*_*i*_ for all *i* is transformed into

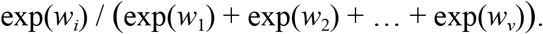

Below we only consider paths that start in a source and end in a sink. Note that the maximum-weight path in *G* is also the maximum-weight path in the transformed graph *G’*.

The adjacency matrix of the transformed graph *G’* can be viewed as a transition matrix for a Markov Chain *Chain*. There is a bijection between the set of paths in *G’* and trajectories of the *Chain*. Thus, the problem of finding a path with the maximum weight in *G* via dynamic programming is equivalent to finding the most likely trajectory of *Chain*.

Since both the standard alignment graph and the modified graph that is used by TandemAligner (defined in subsection “Anchor-based rare-alignment”) are weighted directed acyclic graphs, they can be viewed as Markov Chains. From this probabilistic perspective, a pair of strings *S* and *T generates* an alignment *Alignment* between *S* and *T* with probabilities that are defined via the chosen scoring function. Such a view establishes a duality between combinatorial and probabilistic formulation for alignment.

### Connection between evolution and the alignment problem

Let us assume that there is an evolutionary model of transformations on a string that include *substitutions, deletions* and *insertions* (other transformations such as inversions can be potentially included, but, for simplicity, we exclude them). For simplicity, we will assume that evolution goes backward and then forward in time and consider a string *Descendant* arising from a string *Ancestor* after applying a series of transformations *t*. We define a function *T* that takes as an input a string and a series of transformations and outputs a gapped string where gaps correspond to deletions that are introduced by transformations into the input string — *a*_*D*_ = *T*(*Ancestor, t*). We define a function *R* that removes gap-symbols from a string: *R*(*a*_*D*_) = *Descendant*. We denoteㄱ*t —* a “backward” series of transformations to *t* and denote *a*_*A*_ = *T*(*Descendant*, ㄱ*t), R*(*a*_*A*_) = *Ancestor*. To summarize,

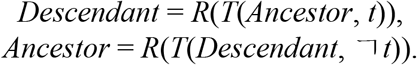

We assume that there is a distribution over transformations with a density *p*(*t*; *Ancestor*). We would like to compute the maximum likelihood alignment (*x*_*A*_,*x*_*D*_) *=Alignment*(*Ancestor, Descendant*) as the mode of the following distribution

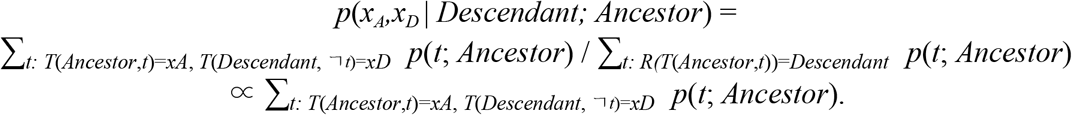

The proportion operation is valid since the denominator does not depend on (*x*_*A*_,*x*_*D*_). The mode of this distribution is the maximum likelihood estimate of *TrueAlignment* denoted as *MLEAlignment*. However, in practice, we do not have access to *p*(*t*; *Ancestor*) (or even the string *Ancestor*)! We attempt to approximate this distribution by defining a Markov Chain (and hence the dynamic acyclic graph) *p*_*MC*_(*Alignment* | *Ancestor, Descendant*; *Scoring*) that depends on the scoring function *Scoring*. This function might either depend on these sequences *Ancestor* and *Descendant* (like the anchor-based alignment that we introduced) or not depend on them (like the traditional alignment approach). We denote the maximum likelihood estimate of *p*_*MC*_ as *MCAlignment*(*Scoring*) to emphasize that this estimate depends on the function *Scoring*.

Finally, the question is how similar is *MCAlignment*(*Scoring*) to *MLEAlignment* (or even *TrueAlignment*)? The answer seems to depend on the underlying distribution *p*(*t*; *Ancestor*). For ETRs, and, in particular, for centromeres, *MCAlignment*(*TandemAlignerScoring*) seems to be closer to *MLEAlignment* than *MCAlignment*(StandardScoring) (see Supplemental Notes “Benchmarking standard and rare-alignments on simulated centromeres”, “Benchmarking standard and rare-alignments on non-repetitive strings”, and “Summary of differences between standard alignment and rare-alignment”).

The emergence of “complete genomics” allows us to revisit previously introduced evolutionary models of centromeres (G. P. Smith 1976) and define alignment scoring schemes that provide better approximations of these evolutionary models. In turn, more accurate alignment scoring schemes provide hypotheses about the evolutionary structure of studied sequences. TandemAligner proposes a simple and conservative scheme that seems to be a more accurate reflection of evolutionary models of ETRs and uses the anchors as the guiding light for the alignment. Future studies should concentrate on this feedback-loop between evolutionary model and alignment scoring scheme.

